# Evolution of cross-tolerance in *Drosophila melanogaster* as a result of increased resistance to cold stress

**DOI:** 10.1101/2020.04.19.047746

**Authors:** Karan Singh, Manas Arun Samant, Nagaraj Guru Prasad

## Abstract

Cold stress is a critical environmental challenge that affects an organism’s fitness-related traits. In *Drosophila*, increased resistance to specific environmental stress may lead to increased resistance to other kinds of stress. In the present study, we aimed to understand whether increased cold stress resistance in *Drosophila melanogaster* can facilitate their ability to tolerate other environmental stresses. For the current study, we used successfully selected replicate populations of *D. melanogaster* against cold shock and their control population. These selected populations have evolved several reproductive traits, including increased egg viability, mating frequency, male mating ability, ability to sire progenies, and faster recovery for mating latency under cold shock conditions. In the present work, we investigated egg viability and mating frequency with and without heat and cold shock conditions in the selected and their control populations. We also examined resistance to cold shock, heat shock, desiccation, starvation, and survival post-challenge with *Staphylococcus succinus* subsp. *succinus* PK-1 in the selected and their control populations.

After cold-shock treatment, we found a 1.25 times increase in egg viability and a 1.57 times increase in mating frequency in the selected populations compared to control populations. Moreover, more males (0.87 times) and females (1.66 times) of the selected populations survived under cold shock conditions relative to their controls. After being subjected to heat shock, the selected population’s egg viability and mating frequency increased by 0.30 times and 0.57 times, respectively, compared to control populations. Additionally, more selected males (0.31 times) and females (0.98 times) survived under heat shock conditions compared to the control populations. Desiccation resistance slightly increased in the females of the selected populations relative to their control, but we observed no change in the case of males. Starvation resistance decreased in males and females of the selected populations compared to their controls.

Our findings suggest that the increased resistance to cold shock correlates with increased tolerance to heat stress, but this evolved resistance comes at a cost, with decreased tolerance to starvation.

## Introduction

Empirical studies on *Drosophila* revealed that the evolution of increased resistance to specific environmental stress may confer an advantage or a disadvantage on their ability to tolerate other types of environmental stresses^1,2,3^. A number of studies confirmed the relationship between multiple stress conditions^1,3,4,5,6,7^. For instance, heat tolerance and resistance to starvation correlate with desiccation resistance and cold tolerance independently^1,3,4,5,6,7^. Insects experiencing transient environmental stress for a single generation are protected from other environmental stresses. For example, exposing *D. melanogaster* to mild cold stress can increase their ability to tolerate heat stress^8^. Cold hardening in *D. melanogaster* confers protection against heat stress^9^. Le Bourg’s^8^ study reported that subjection of *D. melanogaster* to mild starvation followed by a 2-6 h delay to cold stress, could facilitate their ability to resist cold stress. Mild desiccation experience in Springtail can increase their ability to tolerate cold stress^10^. Similarly, *Folsomia candida* house flies experiencing an anoxic condition at 27°C show greater tolerance when exposed to -7°C^7,10,11,12,13^.

A number of laboratory selection studies against different stresses reported cross-tolerance between them^4,7,14,15,16^, such as increased heat tolerance in lines selected for increased resistance to desiccation or heat knockdown time^4, 7^. Similarly, increased cold shock tolerance was observed in lines selected for resistance to chill-coma recovery, heat shock, or desiccation^7^. Additionally, a positive correlation in desiccation resistance was observed between lines selected for chill-coma recovery and cold shock resistance^15,16^. An additional unique mechanism required to adapt to a specific environmental stress might conflict with a mechanism required to adapt to other stresses, thereby leading to trade-offs across stress resistance traits^17,18^. For example, Hoffmann et al.^19^ documented that flies selected for increased starvation resistance had decreased resistance to cold stress. In contrast, those lines selected for increased cold resistance show reduced starvation resistance^2^. However, none of the studies investigated how the evolution of cold stress resistance in the context of egg viability and mating frequency post cold shock can impart cross-resistance with other stresses.

In this study, we aimed to assess the cross-tolerance in the context of egg viability and mating frequency with and without cold and heat shock conditions in selected and control populations of *D. melanogaster*. We also investigated resistance to cold shock, heat shock desiccation, starvation, and survival post-challenge with *Staphylococcus succinus* subsp. *succinus* PK-1 (*Ss*) in the selected and control populations. Our study populations consisted of 5 selected and 5 control populations of *D. melanogaster*, and experiments were conducted over 57-70 generations of selection.

## Results

### Experiment 1: Effect of cold shock on egg viability and mating frequency

To assess the effect of the selection regime in the context of egg viability and mating frequency, we measured the egg viability with and without cold shock conditions in the Freeze Shock Selected populations derived from the BRB populations (FSB or selected) and Freeze Shock Control populations derived from the BRB populations (FCB or control). We did not find a significant effect of selection on egg viability data of 0-6 h post-exposure to cold shock and no shock conditions. However, treatment had a significant effect on egg viability with a more than 90% reduction in egg viability post cold shock compared to no shock treatment (**Fig. 1A; Table S1A)**. We did not observe a significant interaction between selection and treatment **(Table S1A)**. For 24-30 h post-exposure, we found a significant effect of selection, treatment, and selection × treatment interaction on the egg viability **(Table S1B)**. The egg viability of 24-30 h post cold shock exposure of the FSB populations increased by 1.25 times compared to the FCB populations (**Fig. 1B)**. An increase in egg viability could result from an increase in mating frequency in the FSB populations compared to the FCB populations. Therefore, we also measured the mating frequency from the same set up of the experiment that was used to assess egg viability. We observed that the selection regime had a significant effect on mating frequency (**Fig. 1C)**. We also observed that treatment had a significant effect on mating frequency, indicating that cold shocked populations had more matings compared to unshocked populations. We also found a significant effect of selection × treatment interaction on mating frequency (**Table S1C**). Multiple comparisons using Tukey’s HSD test confirmed that post cold shock FSB populations had a 1.57 increase in mating frequency compared to FCB populations (**Fig. 1C)**. Unshocked FSB populations had roughly a 7% increase in mating frequency compared to unshocked FCB populations (**Fig. 1C)**. This result suggests that the basal rate of mating frequency also increased in the FSB populations (**Fig. 1C**). Overall, we show that evolved responses to cold shock in the context of egg viability and mating frequency persist across many generations of selection. These results are in agreement with previous reports^22,24^.

**Figure 1.**
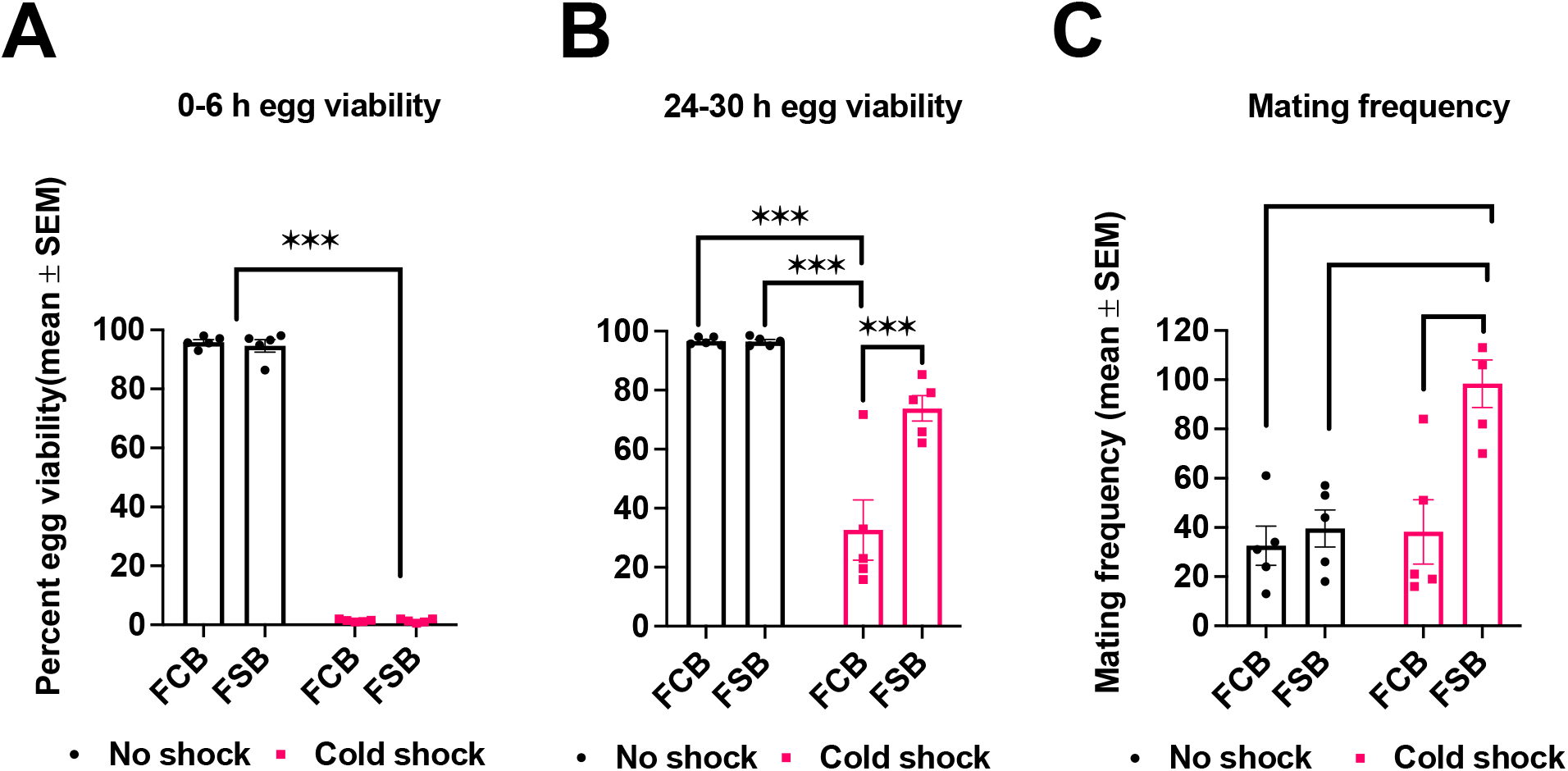
Heightened egg viability and mating frequency in the selected populations post cold shock. (**A** and **B**) Egg viability measurement for each replicate population of FSB and FCB at 0-6 h (**A**) and 24-30 h (**B**) post cold shock and no shock. n = 1 plate (200 eggs/plate) for each FSB and FCB populations and for each treatment condition. **(C)** The number of mating pairs observed every 30 min interval during 36 h post cold shock from each replicate populatio of FSB and FCB. Data are represented as summed of the matings observed across 36 h. n = 5 cages (100 mating pairs of males and females) for FSB and FCB populations for cold shock and no shock treatments. Data are represented as mean ± SEM. **p < 0.01, ***p < 0.001.

### Experiment 2: Effect of heat shock on egg viability and mating frequency

To examine cross-tolerance, we measured egg viability and mating frequency in FSB and FCB populations under heat shock and no shock conditions. For 0-6 h post-exposure to heat shock and no shock treatment, we did not find a significant effect of selection and selection × treatment interaction in egg viability (**Table S2B)**. However, we observed a significant effect of treatment (**Table S2B)**, with heat shocked populations having extremely low egg viability (∼10%) compared to unshocked populations (**Fig. 2A**). For 24-30 h post-exposure to heat shock and no shock treatment, we found a significant effect of selection, treatment, and selection × treatment interaction in egg viability (**Table S2B**). At 24-30 h post heat shock, egg viability was reduced by 34% in the FCB and 43% in the FSB populations compared to the unshocked FCB and FSB populations, respectively. Multiple comparisons using Tukey’s HSD test confirmed that the post heat shock FSB populations had ∼27% more egg viability than the FCB populations (**Fig. 2B**).

**Figure 2.**
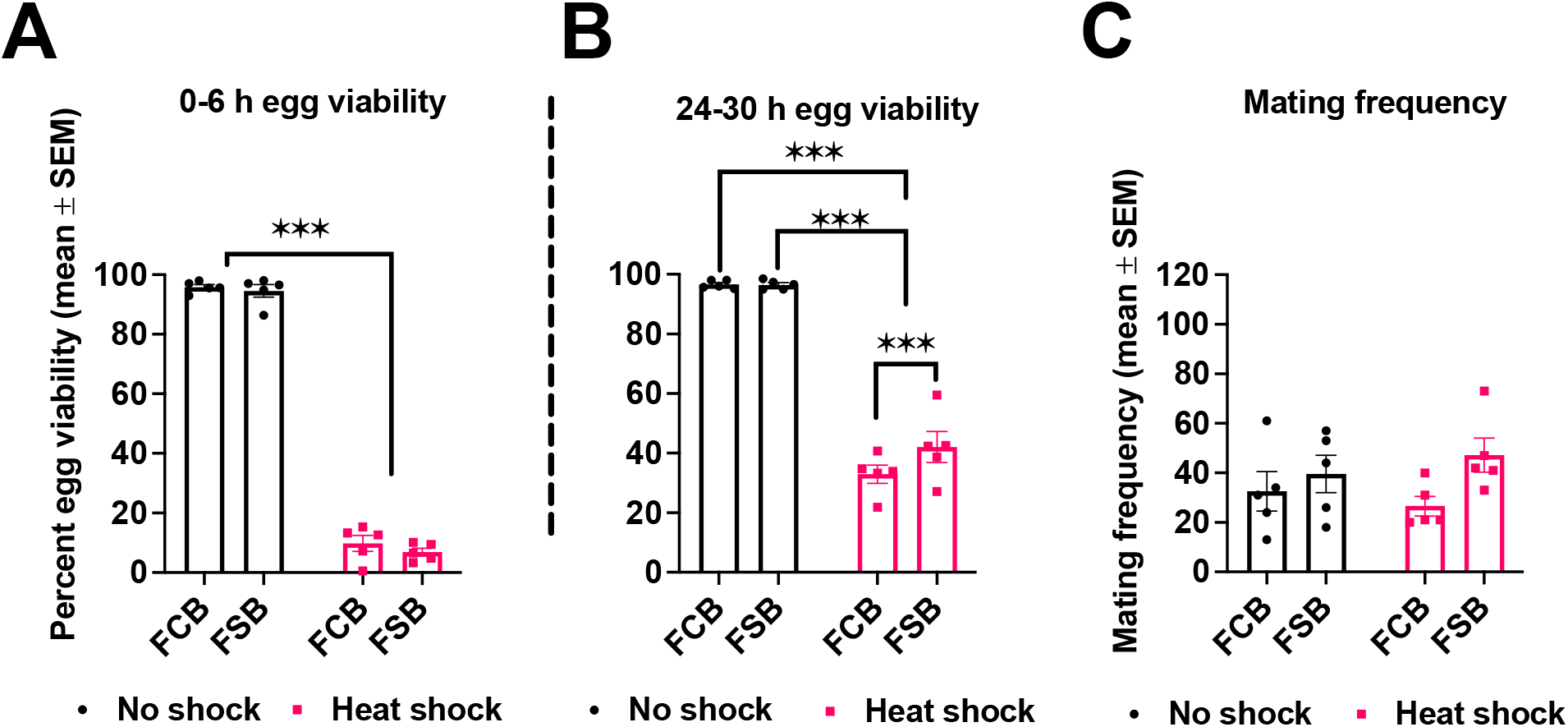
Increased egg viability and mating frequency in the selected populations. (**A** and **B**) Egg viability measurement for each replicate population of FSB and FCB at 0-6 h (**A**) and 24-30 h (**B**) post heat shock and no shock treatment. n = 1 plate (200 eggs/plate) for each replicate population of FSB and FCB, and for heat shock and no shock treatments. (**C**) Post heat shock, the mating frequency was monitored every 30 min intervals for 36 h. n = 5 cages (100 males and females/cage) for selection regime and treatment group. Data are represented as mean ± SEM. ***p < 0.001.

We also measured mating frequency with and without heat shock conditions in the FSB and FCB populations. We observed a significant effect of selection on mating frequency (**Table S2C**). The FSB populations had a 46% increase in mating frequency compared to the FCB populations (**Fig. 2C**). We did not notice any significant effect of treatment, and selection × treatment interaction on mating frequency (**Table S2C**).

### Experiment 3A: Effects of cold shock on adults’ survival

To probe the effect of the selection regime on cold tolerance, we exposed 2-3 days old virgin males -and females of the FSB and FCB populations to cold shock for 1h as described in Methods. Twenty-four hours post cold shock, the survival data of males and females was analyzed. We detected a significant effect of selection on males’ survival post cold shock (**Table S3A)**. At 24 h post cold shock, FSB males had a 0.87 times increased survival compared to FCB males **(Fig. 3A)**. We also observed a significant effect of selection on females’ survival 24 h post cold shock **(Table S3B)**, with FSB females having 1.66 times more survival than FCB females (**Fig. 3A)**.

**Figure 3.**
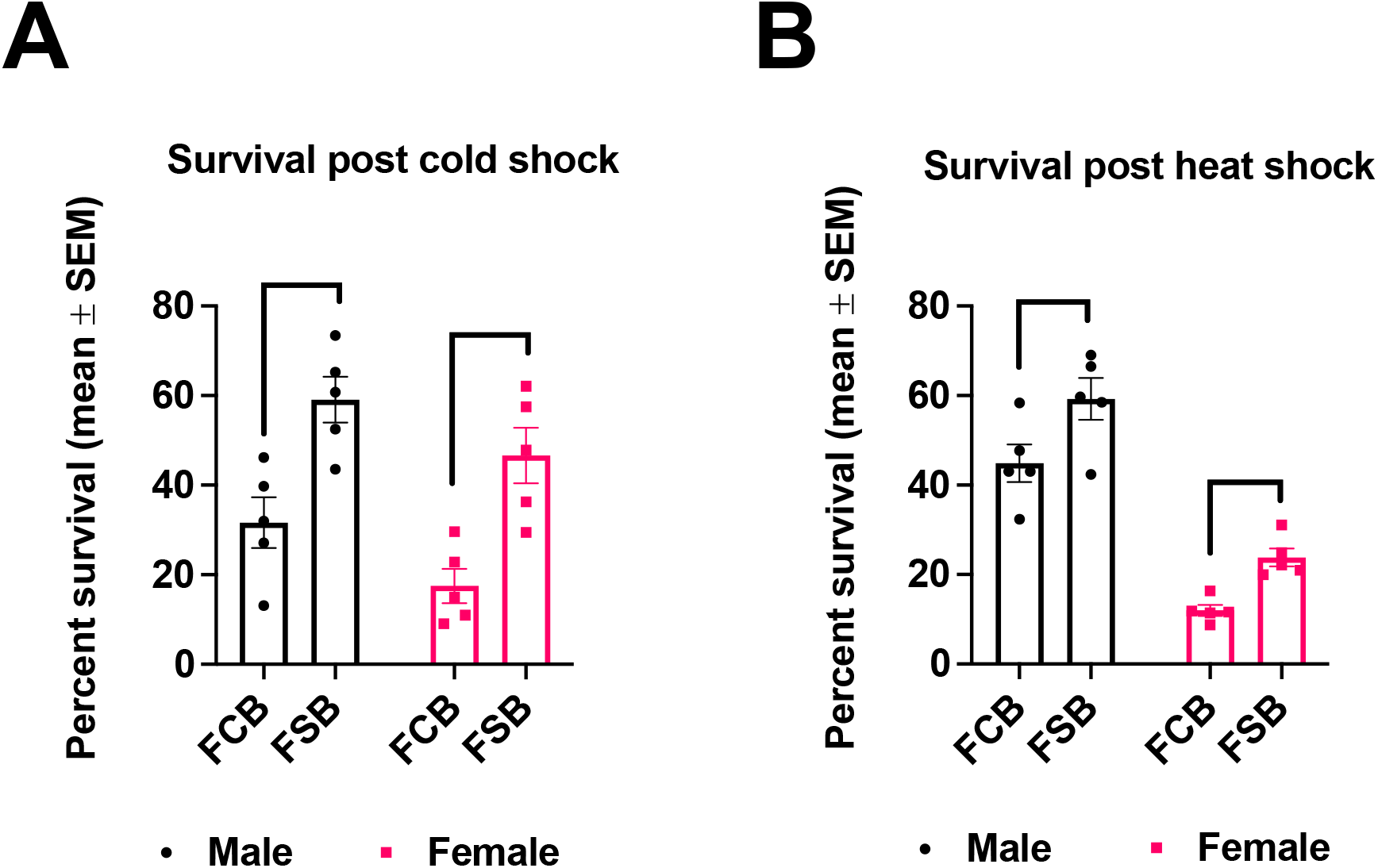
Increased survival in the selected populations post cold and heat shock. **(A)** Percent survival of two-three days old virgin flies of the FSB and FCB populations exposed to cold shock for 1 h as described in Methods. (**B)** Percent survival of two-three days old virgin flies of the FSB and FCB populations exposed to heat shock for 1 has described in Methods. The number of dead flies was counted 24 h after exposure. n = 300 male and female flies at a density of 100 males or females/cage were used for each replicate population of FSB and FCB. Data are represented as mean ± SEM. *p < 0.05, **p < 0.01.

### Experiment 3B: Effects of heat shock on adults’ survival

To assess heat-tolerance in the FSB and FCB populations, virgin male and female flies of the FSB and FCB populations were exposed to heat shock conditions as described in methods. Twenty-four hours after the heat shock, we counted the total number of dead male and female flies from the FSB and FCB populations. We found a significant effect of selection on males’ survival (**Table S3C)**. Twenty-four hours after heat shock, survival of the FSB males had increased by 0.31 times compared to the FCB males (**Fig. 3B)**. Similarly, we also observed a significant effect of selection on females’ survival data 24 h post heat shock (**Table S3D)**. Indicating that 24 h post heat shock, the FSB females had a 0.98 times increase in survival compared with the FCB females (**Fig. 3B)**. These findings suggest that the increased resistance to cold stress in *D. melanogaster* can also confer an ability to resist heat stress.

### Experiment 4: Adult survival post-challenge with Ss

Cold stress is known to activate the immune system in *D. melanogaster*^20^. Therefore, we challenged the male and female flies of the FSB and FCB populations with *Ss*^21^ to determine the effect of the selection regime on adults’ immunity. The survival data of male and female flies post-infection with *Ss* was analyzed using the Cox proportional hazards methods. Our results show no significant difference in survival between FSB and FCB populations **(Fig. 4A** and **B; Table S4)**. We found no significant effect of selection and sex on the survival of male and female flies post-infection with *Ss* (**Table S4)**.

**Figure 4.**
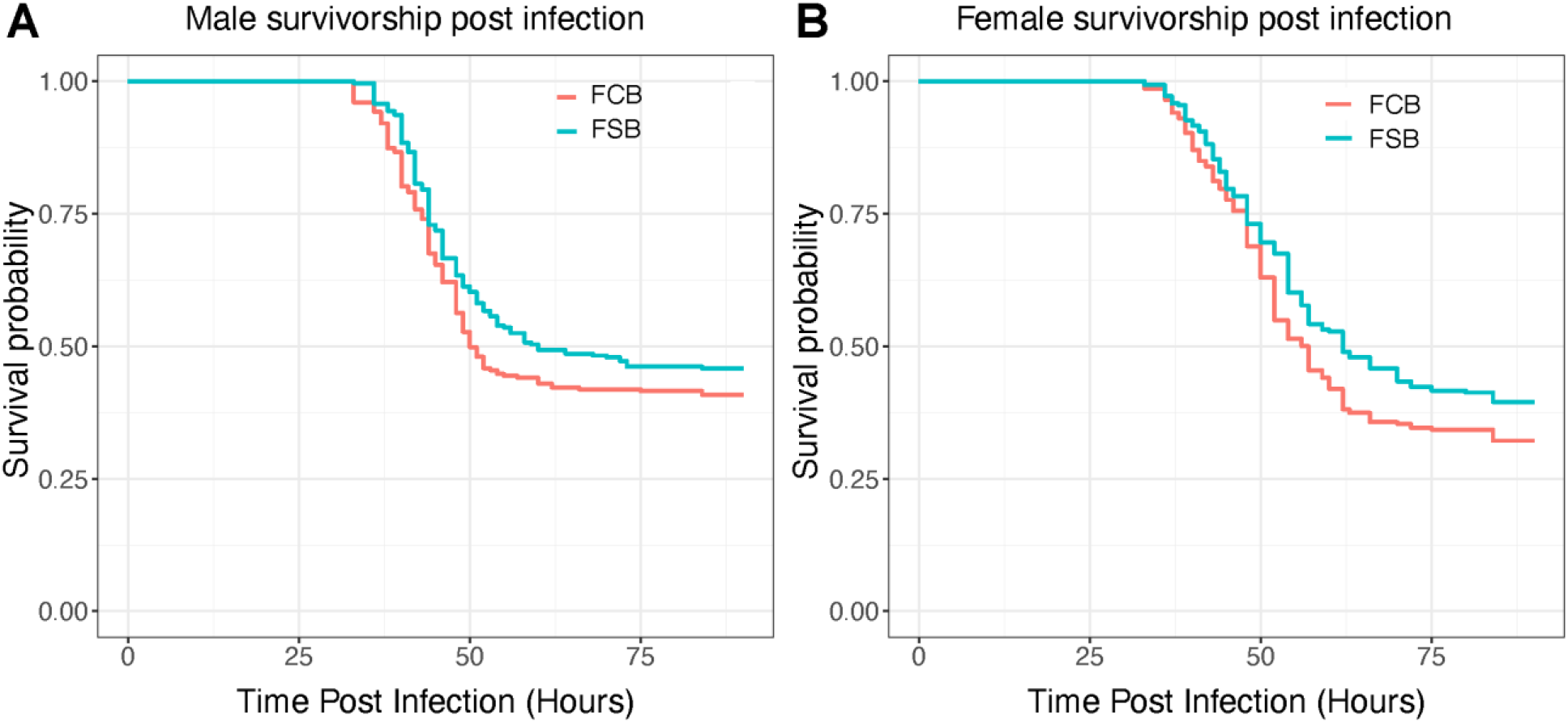
Immune response is not altered in the selected populations. **(A** and **B)** 2-3 days old virgin flies of the FCB or FSB population were challenged with Ss. The number of dead flies were recorded every 3 h intervals for 24h post-infection, and then monitored every 1 h until 90 h post-infection. n = 7 vials of 10 males or females/vial for each of the 5 replicate populations of FSB and FCB. A total of ∼350 males and females flies were assayed from the FSB and FCB populations. Data are represented as survival probability.

### Experiment 5: Desiccation resistance

To determine the effect of the selection regime on desiccation resistance, we exposed male and female flies of the FSB and FCB populations to desiccation conditions. The data were analyzed using Cox-proportional-Hazard model. We found that FSB females had slightly higher desiccation resistance compared to FCB females (**Table S5B; Fig. 5B**). However, in the case of males, we did not find a significant effect of selection (**Table S5A; Fig. 5A**).

**Figure 5.**
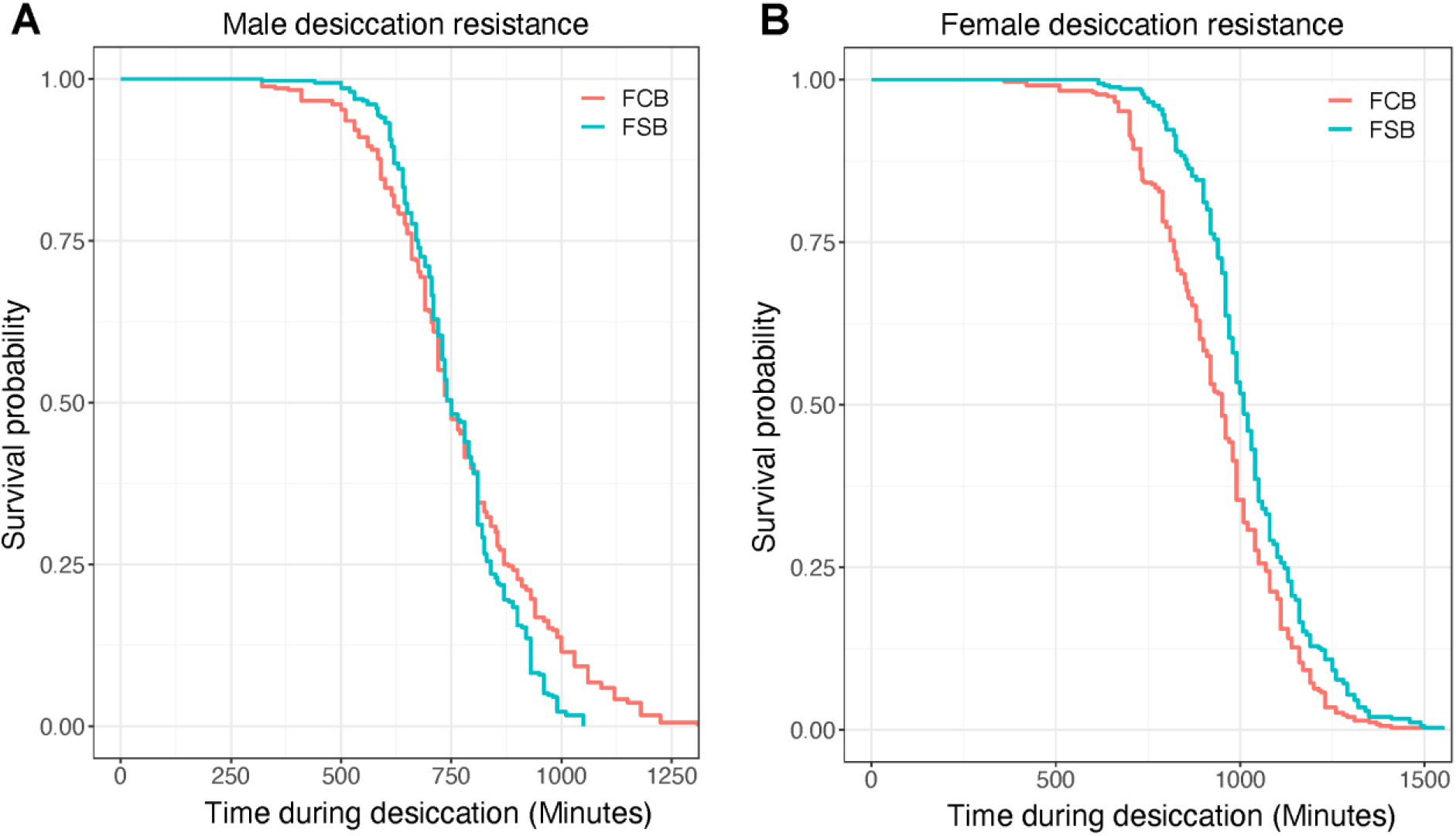
No alteration in desiccation resistance in the selected populations. **(A)** Desiccation resistance of virgin males age of 2-3 days old measured as the number of dead flies every 30 min until all flies were dead. **(B)** Desiccation resistance of virgin females assayed in the same way as the males. n = 7 vials of 10 males or females/vial for each of the 5 replicate populations of FSB or FCB were used for assay. Data are represented as the probability of survival.

### Experiment 6: Starvation resistance

We assayed sex-specific starvation resistance in the FSB and FCB populations after 57 generations of selection, and the survival data was analyzed using the Cox proportional-Hazard model. We observed a significant effect of selection on males’ and females’ starvation resistance (**Table S6A** and **B)**, with the FSB males having significantly reduced starvation resistance compared to FCB males (**Fig. 6A)**. Like FSB males, FSB females also had significantly decreased starvation resistance compared to control females (**Fig. 6B)**. These results suggest that increased cold shock resistance comes with a cost; as a result, there is a trade-off with resistance to starvation.

**Figure 6.**
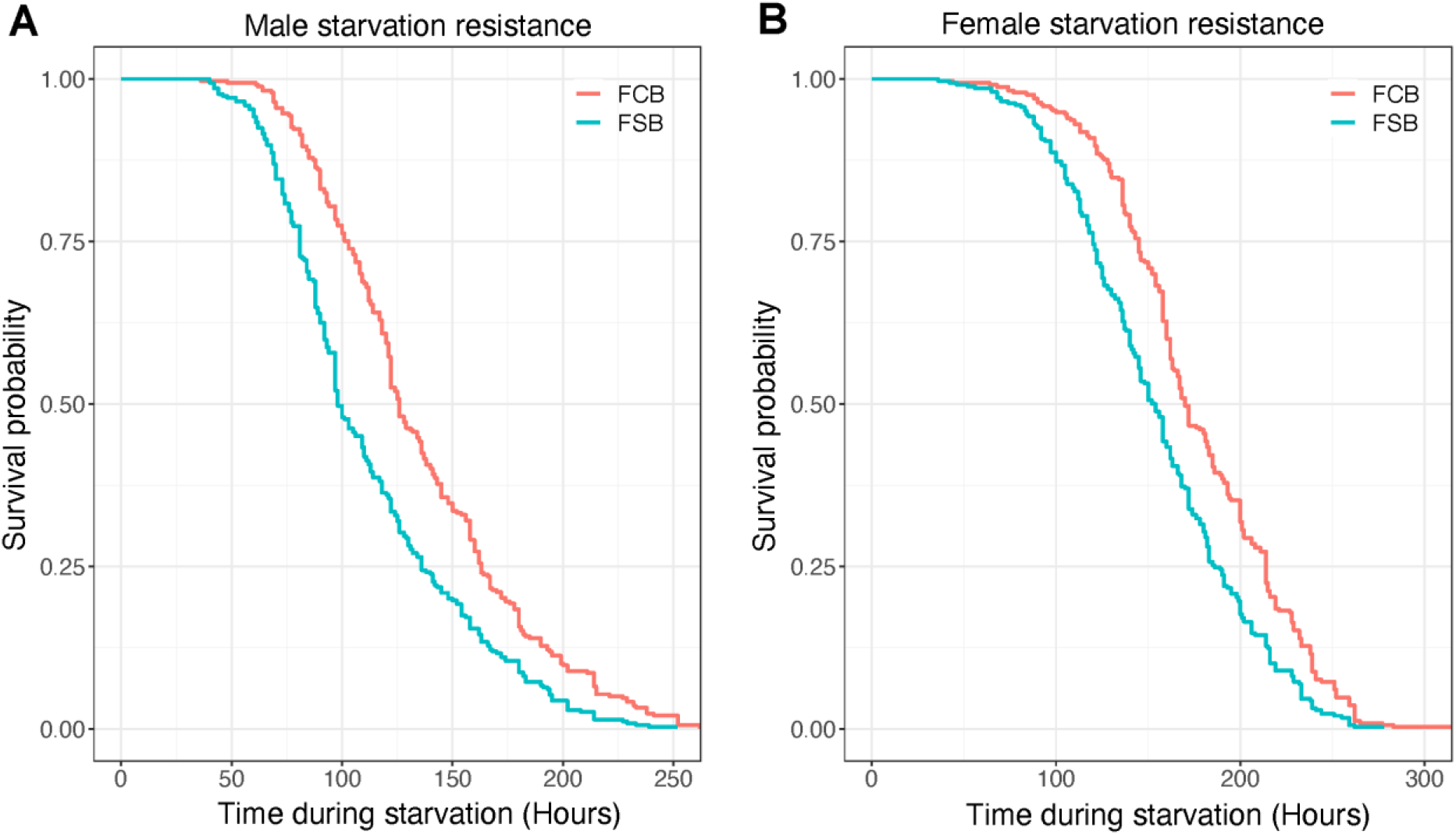
Decreased starvation resistance in the selected populations. **(A)** Starvation resistance measured as the number of dead flies every 4 h until the last fly died. **(B)** Starvation resistance of females, assayed as males. n = 7 vials of 10 males or females/vial for each replicate population of FSB or FCB. Data are represented as a probability of survival.

## Discussion

This study aimed to explore the evolution of cross-tolerance in populations of *D. melanogaster* that were selected for increased resistance to cold shock. These selected populations have evolved multiple reproductive traits, including egg viability, mating frequency, male mating ability, male ability to sire more progenies, mating latency, sperm offense ability, and progeny production^22,23,24^. In our present experimental evolution study, we measured the egg viability and mating frequency with and without heat and cold shock conditions. We investigated the resistance to cold shock, heat shock, starvation, desiccation, and adult survival post-challenge with *Ss* in the selected and control populations. We observed an increased egg viability in the selected populations relative to control populations (**Fig. 1B** and **2B**), as documented in previous studies^22,24^. We also found that the selected populations had a higher mating frequency under cold shock conditions (**Fig. 1C)**, and a slight increase under heat shock conditions **(Fig. 2C)**. After being subjected to cold and heat shock, selected populations had higher adult survival relative to their control populations (**Fig. 3A** and **B)**. Desiccation resistance slightly increased in females of selected populations (**Fig. 5A**), which indicates that selection for an environmental stressor can lead to resistance to other forms of stress. However, in the case of starvation resistance, we found that the selected populations had lower starvation resistance than control populations (**Fig. 6A** and **B**), which suggests that the increased survival of selected populations to cold shock is negatively correlated with starvation resistance. We discussed each of these observations below in more detail.

At 0-6 h post cold shock, we found approximately 95-97.5% reduction in egg viability (**Fig. 1A**). This could be caused by sperm mortality in male seminal vesicles, female seminal receptacles, and spermathecal at below zero temperatures^25,26^. This result is in line with several other studies that have observed reduced egg viability and sterility in insects upon exposure to extreme temperature^10,22,24,27,28^. However, we found greater egg viability in the selected populations compared to control populations 24-30 h after exposure to cold shock (**Fig. 1B**), as it was reported previously^22,24^. There are a number of possible explanations for increased egg viability at 24-30 h post cold shock: (a) The selected populations could be better at protecting their stored sperms/eggs from damage caused by cold shock. For instance, Collett and Jarman^29^ have shown that *D. pseudoobscura* females can store sperm for up to 6 months in cold environments. These stored sperms can be used to fertilize ova in a favorable warm environment. However, this is not the case in our selected populations because a previous report using the same populations showed that females of selected populations relative to their control populations do not store eggs or sperm^22^. (b) The selected populations might have mated more often after heat or cold shock, enhancing egg viability. A number of studies documented that high and low temperatures have an impact on the mating behavior^30,31,32,33,34,35^. However, very few studies have addressed the effect of cold shock on mating behavior^22,23,24^. In the selected populations, mating frequency increased post heat or cold shock compared to control populations (**Fig. 1C** and **2C**). Hence, it is likely that increased mating frequency post heat or cold shock is largely responsible for increase in egg viability. While the pattern of increased mating frequency correlated with increased egg viability post cold shock in the current study as well as in previous reports^22,24^, it is interesting that this pattern is seen to a lesser extent under heat shock conditions. This finding indicates that perhaps some mechanisms underlying resistance to heat and cold stress might be common.

We observed that selected populations had increased survival compared to control populations over 24 h post cold shock (**Fig. 3A**). Our results indicate that the selected populations have evolved the ability to withstand cold stress in terms of increased adult survival along with their ability to maintain higher egg viability after shock at a temperature of -5°C. Multiple laboratory selections studies have shown increased adult survivorship as a correlated response to selection for cold tolerance^2,16,36,37^. Singh et al.^22,24^ documented that the mortality post cold shock was negligible in their selection regime. However, in the present study, mortality post cold shock was substantial (**Fig. 3A**). There are several possible explanations for these contradictory results. First, the populations used in the different studies might have drifted away for a number of generations between the setting of the different experiments. Second, in the current study, the flies were virgins when subjected to cold shock, whereas in the previous study, the flies had already mated by the time they were subjected to cold shock. Third, in the present study, the flies were moved into a fresh food vial soon after eclosion, while in the previous study, the flies remained in the cultured vials (with old, spent food) for two-three days after eclosion. There could be multiple explanations for the superior survivorship of selected populations post cold shock. (a) Chen and Walker^37^ report that cold selected lines have higher glycogen and total proteins relative to control lines. Insects are known to store various sugars in order to tolerate cold temperatures^38,39^. It is possible that the selected populations have similarly altered resource storage in terms of carbohydrates, proteins or lipids. (b) Several studies have shown that there are several heat shock proteins that are expressed both during heat and cold stress. It is quite possible that at least some of these genes are expressed at a higher level in our populations. However, these genes are certainly not among the set that we analyzed for the expression differences in our other study^40^.

Interestingly, the selected populations also showed higher survival post heat shock compared to control populations (**Fig. 3B**). The present literature depicts some disagreement with regards to cross-resistance between cold and heat stress^41^. Anderson et al.^16^ and MacMillan et al.^2^ did not find correlated increase in heat shock resistance in populations of *D. melanogaster* selected for faster chill coma recovery or freeze resistance, respectively. Our results are in agreement with those of Kristensen et al.^42^, who show that cold selected lines of *D. melanogaster* were more heat tolerant and vice versa. Previous studies in *Drosophila* along latitudinal clines suggest that there is a trade-off between heat and cold tolerance^43^. Our results suggest that heat and cold tolerance might be positively correlated in *D. melanogaster*.

In insects, cold stress can cause somatic injury to the gut and malpighian tubules. This can open up a way for the gut flora to enter the haemocoel and thereby cause an infection^44,45,46,47^. Therefore, in our selected populations, immune activity can potentially evolve. However, we noticed that selected populations had a slightly higher survival (**Fig. 4A** and **B**). The difference was not significant between selected and control populations. One possibility is that the immune response is elicited only in response to the gut flora. In *Drosophila*, evolution against a pathogen can be fairly specific, and the host might not have increased immunity against other pathogens^48,49,50^. Thus, in the present assay, where we use *Ss* as the pathogen, the appropriate immune response might not have been elicited.

We found that the desiccation resistance is only slightly increased in females of the selected populations (**Fig. 5B**). Our findings are in line with results from other studies^2,7^, which show that increased resistance to cold shock may lead to increased desiccation resistance as a correlated response. However, populations selected for desiccation resistance do not show increased cold tolerance^15^. There is at least one common factor between cold and desiccation resistance that might explain their correlated evolution. Glycogen is known to act as a cryoprotectant^51,52,53^. Chippindale et al.^53^ showed that selection for increased desiccation resistance leads to increased glycogen content. Thus, increases in glycogen content through selection on cold shock resistance could in principle, lead to the evolution of increased desiccation resistance. Given the sex-specificity of our results, a likely factor that might explain increased desiccation resistance is body size in females. In a previous study, we found that females from selected populations had higher body weight relative to females from control populations^24^.

Starvation resistance decreased in populations selected for increased resistance to cold shock relative to control populations (**Fig. 6A** and **B**). Our findings are similar to those of MacMillan et al.^2^, and Anderson et al.^16^, who found lower starvation resistance in populations of *D. melanogaster* selected for increased resistance to cold shock. Interestingly, Bubliy and Loeschcke^7^ found decreased cold stress tolerance in populations of *D. melanogaster* selected for increased starvation resistance. Thus, across multiple studies, the correlation between starvation resistance and cold stress tolerance seems to be robust.

These findings indicate that the evolution of cold shock resistance provides resistance to heat stress. At the same time, it negatively correlates with starvation resistance. Overall, our results suggest that the increased resistance to cold shock comes at a cost paid by decreased resistance to starvation.

## Methods

### Derivation of selected and control populations

Protocols for the creation and maintenance of the selected have been described previously^22,24^. In short, we established a large, outbred population named BRB in 2010 by mixing 100 male and 100 female flies from each of the 19 isofemale lines, which were originally established in the lab of Prof. Daniel Promislow from 19 inseminated female flies that were captured from the Blue Ridge Mountain, Georgia, USA. In 2010, these 19 isofemale lines were gifted to us. We cultured them for 6 generations on standard laboratory conditions (25°C temperature, 50–60% relative humidity, 12 h shift of light-dark cycles, banana-yeast-jaggery food). We then merged 100 male and female flies from each of the 19 isofemale lines to create a large, outbred population referred to as ‘BRB’. We cultured this BRB population (population size of 2800 flies) for 10 generations under standard laboratory conditions before dividing it into 5 replicate populations known as BRB 1-5. We maintained these BRB 1-5 populations for 35 generations before creating FSB 1-5 and FCB 1-5 from BRB 1-5. Numbers 1-5 denote that both FSB 1 and FCB 1 originated from BRB 1. Therefore, they are more close to each other than FSB 2 and FCB 2 or any other populations; therefore, in the statistical analysis, we considered them as Block 1. Similarly, FSB 2 and FCB 2 were descended from BRB 2; hence, they are more close to each other than FSB 1 and FCB 1 or any other set of populations. Hence, in the statistical analysis, we treated them as block 2 and so on. For additional details, see the below flow chart **(Fig.7)**.

**Figure 7.**
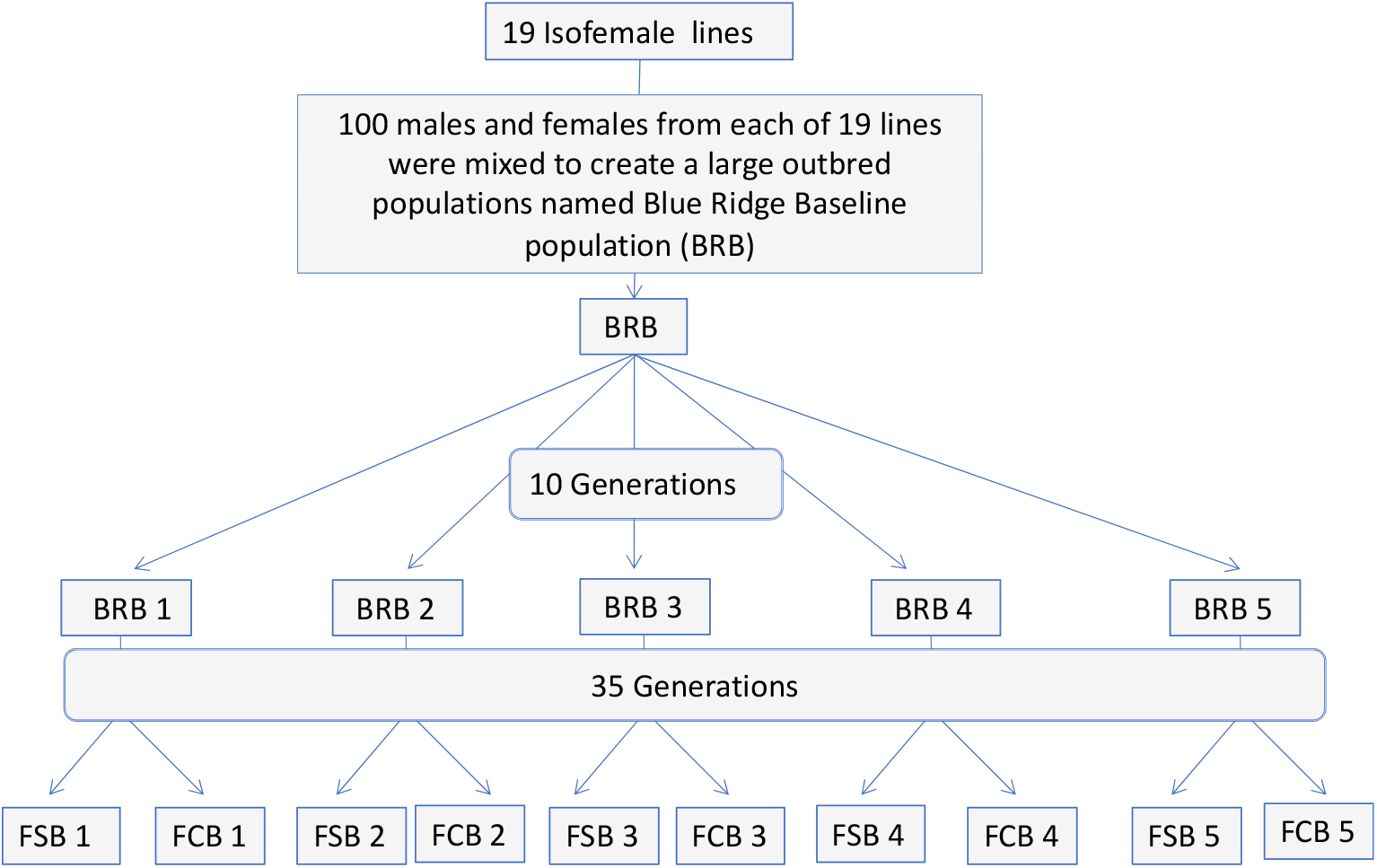
Derivation of the BRB, FSB, and FCB populations.

### Cold shock protocol for selection regime

The FSB 1-5 and FCB 1-5 are large, outbred populations maintained on 13 days discrete generation cycle under standard laboratory conditions. On the 12th day post culturing of eggs, 2-3 days old non-virgin, mated flies at a density of ∼60-70 flies/vial were housed in an empty, clean, dry glass vial (30 mm diameter□×□90 mm length) and a cotton plug was pushed inside the vial so that flies were in a 1/3 area of a vial. Twenty such vials were made for each replicate populations of FSB 1-5 and FCB 1-5. These vials of FSB 1-5 populations were held in a water-ice-salt slurry maintained at -5°C, whereas vials containing flies of the FCB 1-5 populations were kept in a water-bath maintained at 25°C for 1 h. Soon after the treatment, flies were transferred to Plexiglas cages with standard food plate at a density of 1200-1400 flies per cage for each populations of FSB 1-5 and FCB 1-5. To initiate the next generation, 24 h after cold shock, a fresh food plate was given to each cage of FSB 1-5 and FCB 1-5 populations for 18 h to collect eggs. For each of the FCB 1-5 populations, 20 vials of ∼70 eggs/vial were collected so that the total number of flies that emerged from these vials were roughly 1200-1400 flies per population. Cold shock treament is known to affects egg viability, and egg viability at 24-30 h after cold shock was ∼35-40%. Therefore, to control the density of flies per vial for FSB 1-5 populations, 20 vials of ∼100 eggs/vial were collected for each replicate population so that the total number of flies eclosed from these vials were ∼1200-1400 files/population. For more information, see below the graphical illustration (**Fig. 8)**.

**Figure 8.**
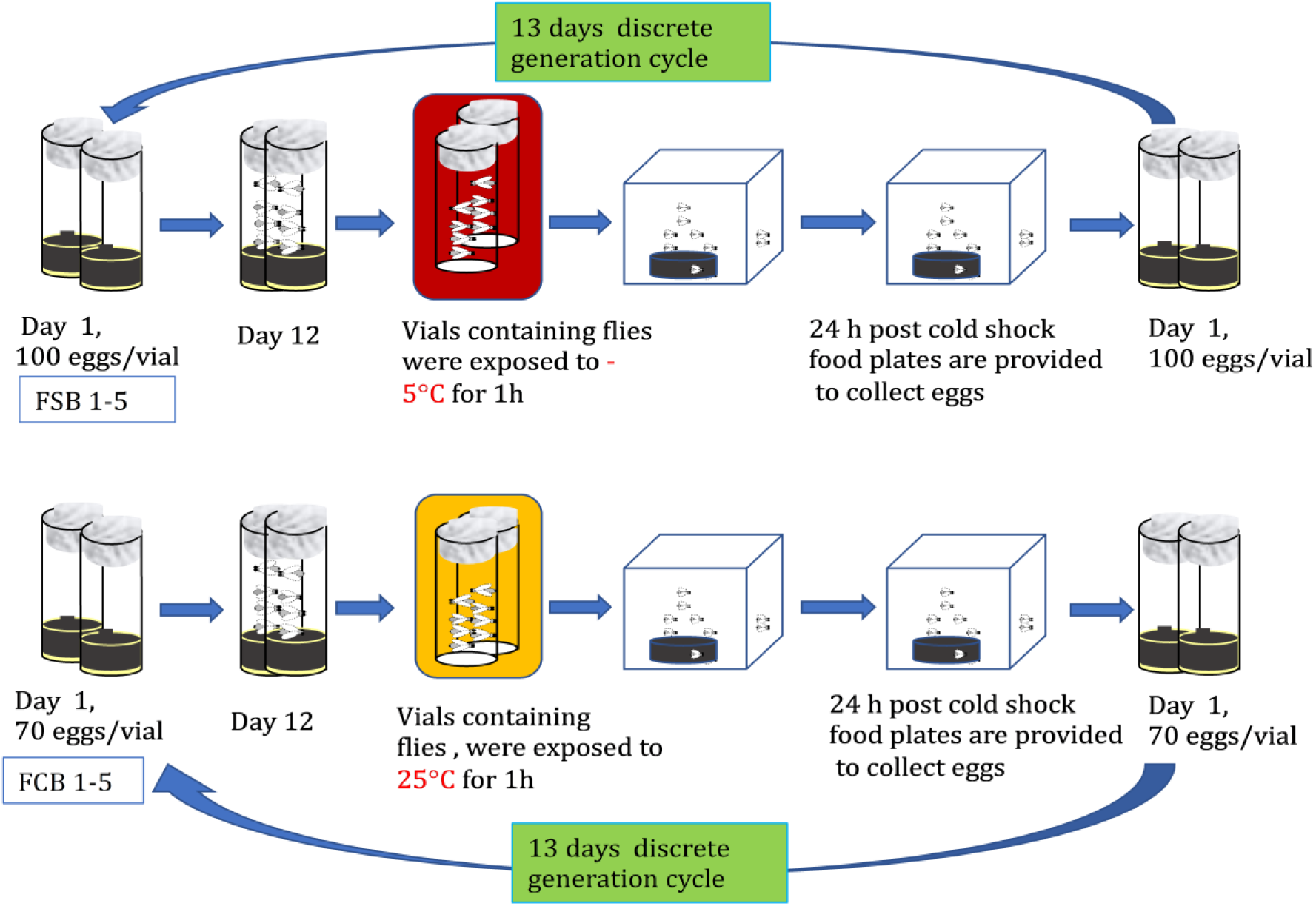
Maintenance of selected and control populations.

### Experimental protocol

#### Standardization of flies

To control over the non-genetic parental effects for selected and control populations ^54,55^, all populations of FSB 1-5 and FCB 1-5 were reared for one generation under common standard laboratory conditions as described below, and we did not impose any selection to FSB 1-5 and FCB 1-5 populations for one generation. This process is known as standardization, and the flies maintained in this manner are known as standardized flies. To standardize the flies, we collected eggs at a density of 70 eggs/vial in a vial with 6 mL of banana-yeast-jaggery food (standard food). We collected 20 such vials from each stock populations of FSB 1-5 and FCB 1-5. On day 12, after culturing eggs, the flies from a specific population were shifted into a Plexiglas cage containing a standard food plate. In order to collect experimental eggs, on day 13 post egg collection, a standard fresh food plate was given to each of the FSB 1-5 and FCB 1-5 populations, and the flies were allowed to oviposit for 6 h. To raise experimental flies from each of the FSB 1-5 and FCB 1-5 populations, eggs were cultured at a density of 70 eggs/vial in a vial with 6 mL of standard food as specified in below experiments. All experiments in the present study were conducted over 57 to 70 generations of the selection.

#### Experiment 1-2: Effect of heat shock or cold shock on the mating frequency and egg viability

To probe the effect of the selection regime on egg viability and mating frequency under heat, cold, and no shock conditions, we collected eggs at a density of 70 eggs/vial in a vial containing 6 mL food. Twelve such vials were set up from each of the standardized flies of FSB 1-5 and FCB 1-5 populations. On day 12 post culturing of eggs, non-virgin flies were grouped at a density of 25 males and females/vial (n = 4 vials/block/treatment, therefore, total 60 vials for FSB and FCB populations were set up) and 20 vials were randomly assigned to one of the following three treatments for FSB and FCB : **(a) Cold-shock:** non virgin flies were exposed to - 5°C temperature for 1 h, as defined above. Post cold shock, flies were quickly transferred to a Plexiglas cage containing a fresh food plate at a density of 100 mating pairs per cage/block (n = 5 cages for the FSB 1-5 and FCB 1-5 populations). **(b) Heat-shock:** non virgin flies were exposed to 37.5°C for 1 h. Post heat shock, flies were quickly transferred to a Plexiglas cage containing a fresh food plate at a density of 100 mating pairs per cage/block (n = 5 cages for the FSB 1-5 and FCB 1-5 populations). **(c) No-shock:** non virgin flies were exposed to 25°C for 1 h (as described above) and subsequently flies were transferred to a Plexiglas cage at a density of 100 mating pair per Plexiglas cage (n = 5 cages for FSB 1-5 and FCB 1-5 populations). We used the same data of no shock treatment to analyze cold shock vs no shock, and heat shock vs no shock treatments. After the cold shock, heat shock, and no shock treatment, we measured the egg viability at 0-6 h and 24-30 h post-exposure from FSB and FCB population. We selected these two periods; (a) 0-6 h and (b) 24-30 h for egg viability measurements (a) because 0-6 h would show the immediate effect of treatment, and (b) 24-30 h represents the effect of selection in their routine maintenance cycle as eggs are collected from 24-42 h after treatment to start the next generation. More detail of the experimental design is shown in the illustration (**Fig. 9)**.

**Figure 9.**
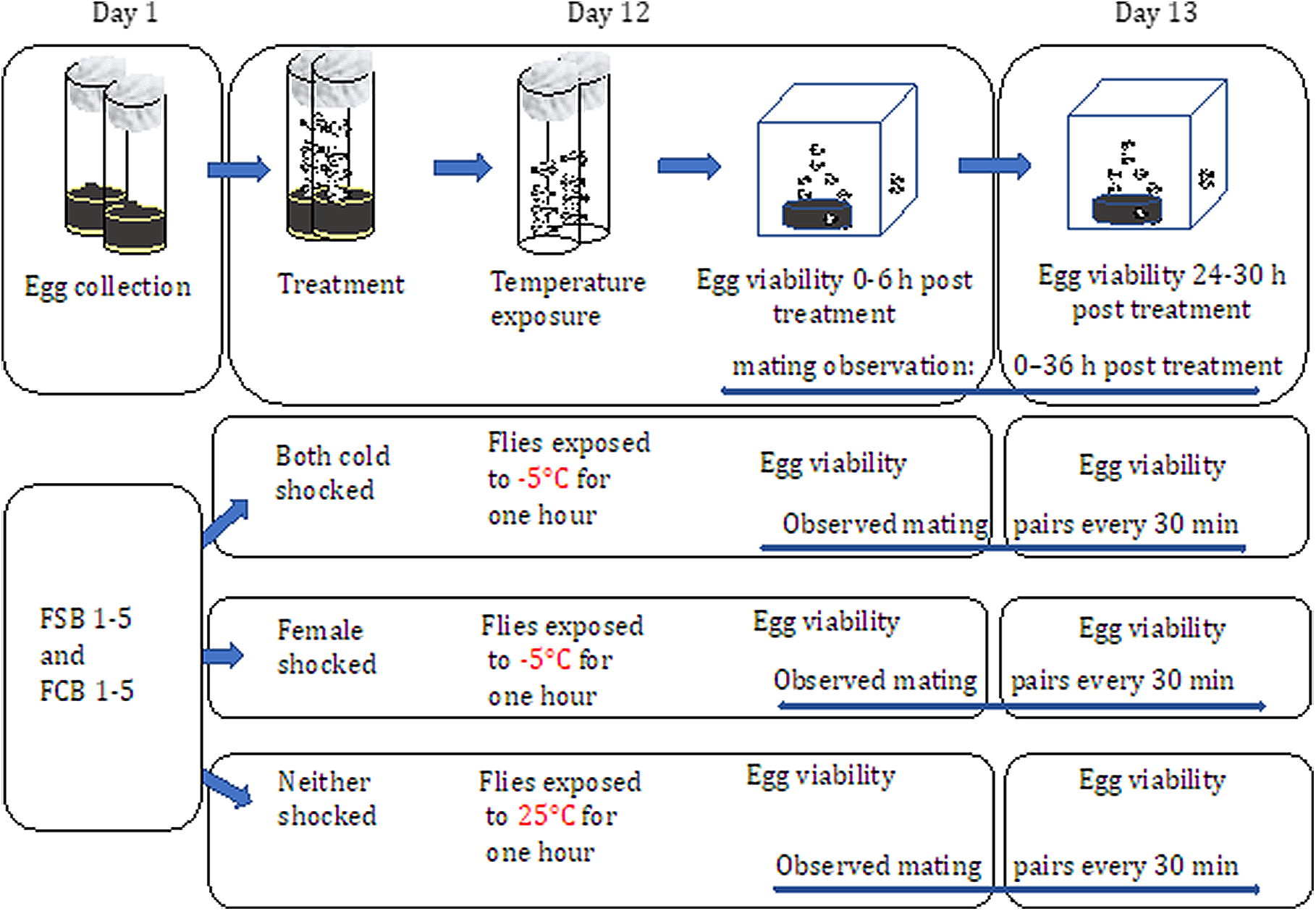
Assay of egg viability and mating frequency.

To measure egg viability, at 0-6 h and 24-30 h post cold shock, heat shock, and no shock treatment to FSB and FCB populations, fresh standard food plates were kept in the FSB and FCB cages for flies to lay eggs for 6 h. After that, eggs at a density of ∼200 eggs/plate were transferred to a Petri plate containing 1.2% agar from the standard food plate that was given to FSB or FCB populations. Following this, the plates were incubated at standard laboratory conditions as described above for 48 h. The number of hatched eggs were counted to compute the percent egg viability using a formula given in the bracket [Percent egg viability = (Number of hatched eggs/ Total number of eggs) × 100]. We also determined the mating frequency from **Experiment 1**. Post cold shock, heat shock, and no shock treatment, flies of the FSB and FCB populations were quickly shifted to Plexiglas cages at a density of 100 pairs of male and female flies per cage. n = 1 cage at a density of 100 male and female flies/cage for each FSB 1-5 and FCB 1-5 populations was established. We scanned all cages every 30 minutes intervals over 36 h post treatment to monitor mating pairs using the same protocol that was reported previously^22,24^. We then summed the number of mating pairs per cage across all 36 hours of observations to estimate the total number of matings. More detail of the experimental design is shown in the illustration (**Fig. 9)**.

#### Experiment 3A: Effect of cold shock on adults’ survival

To investigate the effect of the selection regime on adult survival post cold shock, we assayed survival under cold shock conditions after 63 generations of selection. Eggs were cultured in a standard food vial from the standardized flies of FSB 1-5 and FCB 1-5 populations. Twenty-five vials (70 eggs/vial) were set up for each replicate populations of the FSB and FCB. On the 9-10th day post culturing of eggs, virgin male and female flies were collected from the peak of eclosion using light CO_2_ anesthesia at a density of 10 male or female flies/vial with 2 mL of standard food. Two-three days old virgin male or female flies were segregated in an empty glass vial at a density of 50 flies per vial, and a cotton plug was pushed deep into a vial so that flies were restricted to a 1/3 area of the vial. Female flies were exposed to -5°C for 1 h, and male flies were exposed to -5.6°C for 1 h (we used -5.6°C for males to get at least 50% mortality post cold shock). Post cold shock, male and female flies were immediately transferred to the Plexiglas cage at a density of 100 male or female flies per cage. Twenty-four hours post cold shock, we monitored adult survival. We selected this time point because 24 h post cold shock is the time when fresh food plates are given to FSB 1-5 and FCB 1-5 populations to collect eggs to initiate the next generation in their regular maintenance cycle and hence, measurement at this time is directly relevant to the fitness of the flies. Twenty-four hours post cold shock, dead flies (if any) were aspirated out of the cage and counted. The mean percent survival for each cage of the FSB and FCB populations were computed and used as the unit of analysis. The data were analyzed using a two-factor mixed model ANOVA. n = 3 replicate cages at a density of 100 male or female flies/cage for each replicate population of FSB 1-5 and FCB 1-5. Therefore, about 1500 male and female flies were used in the assay from the FSB and FCB populations. More detail of the experimental design, see the illustration (**Fig. 10)**.

**Figure 10.**
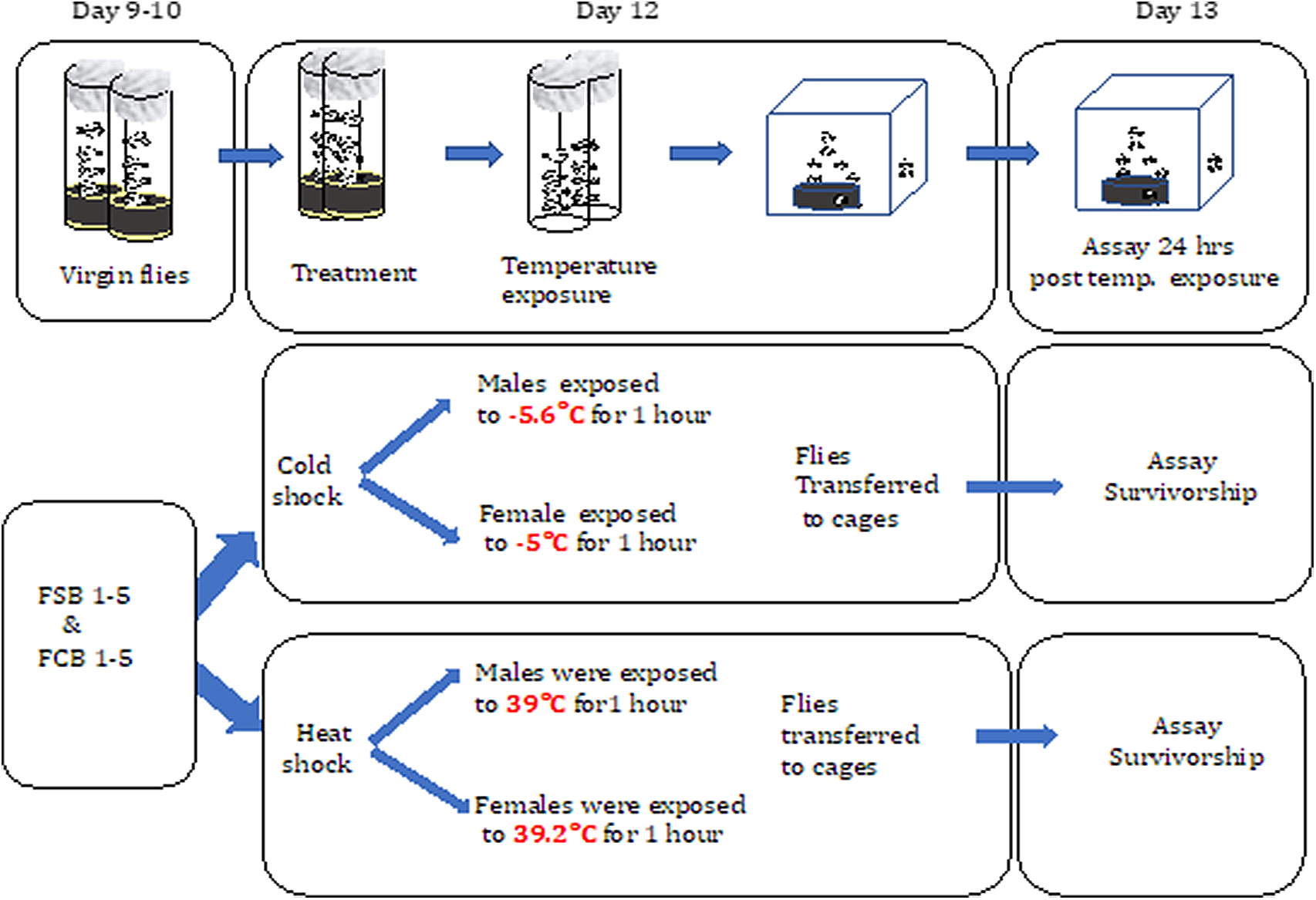
Adults’ survival post heat and cold shock.

#### Experiment 3B: High temperature tolerance

To test heat shock tolerance in the FSB and FCB populations, we assayed adults’ survival post heat shock after 63 generations of selection. The experimental design is the same as it was described for **Experiment 3A**. Both male and female flies of FSB and FCB populations were handled identically as described for **Experiment 3A** except for the temperature treatment. For heat shock treatment, males were exposed to 38.9°C temperature for 1 h in a water-bath, whereas females were exposed to 39.2°C temperature for 1 h in a water-bath. Twenty-four hours post heat shock, dead flies (if any) were aspirated out of the cage, and the percent survival of male and female flies was computed. Statistical analysis was done using a two-factor mixed model ANOVA. n = 3 replicate cages at density of 100 male or female flies/cage were assayed for each replicate population of FSB 1-5 and FCB 1-5. Therefore, a total of about 1500 male and female flies from FSB and FCB population were assayed. More detail of the experimental design is shown in the illustration (**Fig. 10)**.

#### Experiment 4: Adult survival post-challenge with a Ss

After 70 generations of selection, experimental flies were generated as described above for **Experiment 3A**. On day 12 post egg collections, roughly 2-3 days old, 55-60 virgin male and female flies from each replicate population of FSB 1-5 and FCB 1-5 were infected by pricking the lateral thorax with a Minutien pin (0.1 mm, Fine Science Tools, Foster City, CA, USA) that was dipped in the slurry of *Ss* which had OD_600nm_ 2^21^. For sham control, the pin was dipped in 10 mM MgSO4 prior to pricking the flies’ lateral thorax. Dead flies in each vial were monitored every 3 h intervals for 30 h post-infection. After that, vials were observed every hour until 90 h post-infection. n = ∼285-300 male and female flies from the FSB and FCB populations were used in the assay. The survival data was analyzed using a Cox-proportional-Hazard model.

#### Experiment 5: Desiccation resistance

Sex-specific desiccation resistance assay was performed for each FSB (1-5) and FCB (1-5) populations. After 57 generations of selection, experimental flies were raised, as described above in **Experiment 3A**. Two-three days old, virgin male or female flies at a density of 10 flies/vial were transferred from food vials to food-less vials containing ∼6g of silica gel (desiccant). The flies were separated from the silica gel by a thin layer of cotton, and the open end of each vial was sealed with Parafilm as described previously^56^. Mortality was monitored every half an hour until the last fly died in a given vial. n = 7 replicate vials at a density of 10 male, or female flies/vial were set up for each replicate population of FSB and FCB; therefore, about 285-300 male and female flies from FSB and FCB populations were assayed. The statistical significance of desiccation resistance was confirmed with the Cox-proportional-Hazard model.

#### Experiment 6: Starvation resistance

To determine cross-tolerance with starvation resistance in the populations selected for increased resistance to cold shoc<u>k, we</u> assayed sex-specific starvation resistance after 57 generations of selection following the previously described protocol in Kwan et al.^56^ with minor modifications. Experimental flies were generated as described in **Experiment 3A**. On day 12 post egg collections, 2-3 days old, 10 virgin male or female flies were transferred from a food vial to a vial containing 1.24% agar. Flies were transferred into a fresh agar vial (1.24%) every alternate day until the last fly died. Mortality was recorded every four hours. n = 7 replicate vials at a density of 10 male, or female flies/vial were set up for each replicate population of FSB and FCB. Therefore, about 285-300 male and female flies from FSB and FCB populations were assayed. The statistical significance of desiccation resistance data were evaluated with the Cox-proportional-Hazard model.

#### Statistical analysis

Egg viability and mating frequency data of **Experiments 1A-C and 2A-C** were analyzed using a three-factor mixed model ANOVA with GLM considering selection regime (FSB vs. FCB), treatment (cold shock vs. no shock) as the fixed factors crossed with block (1-5) as a random factor. Analyses were performed using a GLM with binomial errors and a logit link function (*i*.*e*., logistic regression) using the R package lme4. Details of the analysis are given in the supplementary information file. Adult’s survival of post heat or cold stress data from **Experiments 3A** and **B** were analyzed using a two-factor mixed model ANOVA considering selection regime (FSB vs. FCB) as a fixed factor crossed with random blocks (1-5). Survival data post infection with *Ss* (**Experiment 4**), desiccation resistance (**Experiment 5**), and starvation resistance (**Experiment 6**) were analyzed using Cox proportional-hazard model in R package Coxme StatSoft.

## List of abbreviations

FSB: Freeze shock Selected populations derived from the BRB populations
FCB: Freeze Shock Control populations derived from the BRB populations
BRB: Blue Ridge Baseline populations
ANOVA: Analysis of Variance
h: Hour
g: Gram
mL: Milliliter

## Decelerations

### Ethics approval and consent to participate

Not applicable.

### Consent for publication

Not applicable.

### Availability of data and material

All data files will be available from the DRYAD database, once the manuscript is accepted.

### Funding

This study was supported by the Indian Institute of Science Education and Research Mohali, Govt. of India. The funders had no role in conceptualizations, investigations, design of study, data collection, analysis, decision to publish, and preparation of the manuscript.

### Authors’ contributions

KS: Conceptualized, designed the study, performed the experiments, analyzed the data, visualized the data wrote an original draft, edited and revised the manuscript. MAS: Contributed to performing immunity experiments, visualized the data, and analyzed the data. NGP: Contributed reagents, materials, and analysis tools. All authors have read and approved the manuscript.

## Acknowledgments

Karan Singh thanks to Indian Institute of Science Education and Research Mohali and Ministry of Human Resources Development, Government of India for financial support in the form of Junior and Senior Research Fellowship. Manas A Samant thanks to the Council of Scientific and Industrial Research, Government of India for financial support in the form of Junior and senior research fellowship. We also thank Megha Kothari for her assistance in the laboratory.

## Supplementary Information

**Table S1A.** Immediate effect of cold shock on 0-6 h of egg viability.

|  | Sum Sq | Mean Sq | NumDF | DenDF | F value | Pr(>F) |
| --- | --- | --- | --- | --- | --- | --- |
| Selection (Sel) | 2 | 2 | 1 | 5.2645 | 0.3608 | 0.5730 |
| Treatment (Trt) | 33185 | 33185 | 1 | 6.0548 | 5976.7529 | **2.671e-10** |
| Sel:Trt | 2 | 2 | 1 | 4.5212 | 0.3458 | 0.5846 |

|  | npar | logLik | AIC | LRT | Df | Pr(>Chisq) |
| --- | --- | --- | --- | --- | --- | --- |
| <none> | 8 | -40.977 | 97.954 |  |  |  |
| (1 \| Block) | 7 | -40.977 | 95.954 | 0.000000 | 1 | 0.9999 |
| (1 \| Block:Selection) | 7 | -40.991 | 95.982 | 0.028029 | 1 | 0.8670 |
| (1 \| Block:Treatment) | 7 | -41.040 | 96.079 | 0.125469 | 1 | 0.7232 |

Warning message:
In checkConv(attr(opt, "derivs"), opt\$par, ctrl = control\$checkConv, :Model failed to converge with max|grad| = 0.172747 (tol = 0.002, component 1)

**Table S1B.** Increased egg viability at 24-30 h post cold shock in the selected populations.

885 >anova(fit\_ev\_cs\_24-30 h) 886 Type III Analysis of Variance Table with Satterthwaite's method
|  | Sum Sq | Mean Sq | NumDF | DenDF | F value | Pr(>F) |
| --- | --- | --- | --- | --- | --- | --- |
| Selection (Sel) | 1881.8 | 1881.8 | 1 | 4.0009 | 21.281 | **0.009927** |
| Treatment (Trt) | 4052.7 | 4052.7 | 1 | 4.0051 | 45.831 | **0.002473** |
| Sel:Trt | 2136.2 | 2136.2 | 1 | 4.0002 | 24.158 | **0.007956** |

|  | npar | logLik | AIC | LRT | Df | Pr(>Chisq) |
| --- | --- | --- | --- | --- | --- | --- |
| <none> | 8 | 65.531 | -147.06 |  |  |  |
| (1 \| Block) | 7 | -65.531 | 145.06 | 0.00059 | 1 | 0.9807 |
| (1 \| Block:Selection) | 7 | -65.538 | 145.07 | 0.01371 | 1 | 0.9068 |
| (1 \| Block:Treatment) | 7 | -65.970 | 145.94 | 0.87831 | 1 | 0.3487 |

pairs(emmeans(fit\_ev\_cs\_24, ~Selection\*Treatment))
| contrast | estimate | SE | df | t.ratio | p.value |
| --- | --- | --- | --- | --- | --- |
| FCB cold shock - FSB cold shock | -41.2 | 6.13 | 7.97 | -6.728 | **0.0007** |
| FCB cold shock - FCB no shock | -64.0 | 7.66 | 6.91 | -8.357 | **0.0003** |
| FCB cold shock - FSB no shock | -63.9 | 7.80 | 7.14 | -8.192 | **0.0003** |
| FSB cold shock - FCB no shock | -22.8 | 7.80 | 7.14 | -2.918 | 0.0820 |
| FSB cold shock - FSB no shock | -22.7 | 7.66 | 6.91 | -2.959 | 0.0798 |
| FCB no shock - FSB no shock | 0.1 | 6.13 | 7.97 | 0.016 | 1.0000 |

*p*-values in bold are statistically significant.
Degrees-of-freedom method: kenward-roger
P value adjustment: tukey method for comparing a family of 4 estimates

**Table S1C.** Higher mating frequency in the selected populations.

|  | Sum<br>Sq | Mean<br>Sq | Num<br>DF | Den<br>DF | F value | Pr(>F) |
| --- | --- | --- | --- | --- | --- | --- |
| Selection (Sel) | 292.6 | 292.6 | 1 | 3.9998 | 22.208 | **0.009223** |
| Treatment (Trt) | 390.1 | 390.1 | 1 | 3.9999 | 29.611 | **0.005537** |
| Sel:Trt | 3537.8 | 3537.8 | 1 | 4.0001 | 268.530 | **8.117e-05** |

|  | npars | logLik | AIC | LRT | Df | Pr(>Chisq) |
| --- | --- | --- | --- | --- | --- | --- |
| <none> | 8 | -67.088 | 150.18 |  |  |  |
| (1 \| Block) | 7 | -68.157 | 150.31 | 2.1388 | 1 | 0.143616 |
| (1 \| Block:Selection) | 7 | -70.436 | 154.87 | 6.6979 | 1 | **0.009653** |
| (1 \| Block:Treatment) | 7 | -69.779 | 153.56 | 5.3828 | 1 | **0.020336** |

*p*-values in bold are statistically significant. pairs(emmeans(fit\_mr\_cs, ~Selection\*Treatment))
| **contrast** | **estimate** | **SE** | **df** | **t.ratio** | **p.value** |
| --- | --- | --- | --- | --- | --- |
| FCB cold shock - FSB cold shock | -60.2 | 7.31 | 4.41 | -8.233 | **0.0028** |
| FCB cold shock - FCB no shock | 5.6 | 6.14 | 4.60 | 0.913 | 0.8003 |
| FCB cold shock - FSB no shock | -1.4 | 9.27 | 7.74 | -0.151 | 0.9987 |
| FSB cold shock - FCB no shock | 65.8 | 9.27 | 7.74 | 7.102 | **0.0005** |
| FSB cold shock - FSB no shock | 58.8 | 6.14 | 4.60 | 9.583 | **0.0012** |
| FCB no shock - FSB no shock | -7.0 | 7.31 | 4.41 | -0.957 | 0.7788 |

*p*-values in bold are statistically significant. Degrees-of-freedom method: kenward-roger P value adjustment: tukey method for comparing a family of 4 estimates

**Table S2A:** Immediate effect of heat shock on egg viability of 0-6 h.

|  | **Sum Sq** | **Mean Sq** | **Num DF** | **Den DF** | **F value** | **Pr(>F)** |
| --- | --- | --- | --- | --- | --- | --- |
| Selection (Sel) | 22 | 22 | 1 | 16 | 1.2643 | 0.2774 |
| Treatment (Trt) | 37732 | 37732 | 1 | 16 | 2146.12 | **<2e-16** |
| Sel:Trt | 3 | 3 | 1 | 16 | 0.1989 | 0.6616 |

|  | **npar** | **logLik** | **AIC** | **LRT** | **Df** | **Pr(>Chisq)** |
| --- | --- | --- | --- | --- | --- | --- |
| <none> | 8 | -48.857 | 113.71 |  |  |  |
| (1 \| Block) | 7 | -48.857 | 111.71 | 0 | 1 | 1 |
| (1 \| Block:Selection) | 7 | -48.857 | 111.71 | 0 | 1 | 1 |
| (1 \| Block:Treatment) | 7 | -48.857 | 111.71 | 0 | 1 | 1 |

**Table S2B.** Selected population have heightened egg viability at 24-30 h post heat shock.

|  | **Sum Sq** | **Mean Sq** | **NumDF** | **DenDF** | **F value** | **Pr(>F)** |
| --- | --- | --- | --- | --- | --- | --- |
| Selection (Sel) | 63.00 | 63.00 | 1 | 8.0000 | 30.290 | **0.0005713** |
| Treatment (Trt) | 1186.19 | 1186.19 | 1 | 4.0016 | 570.328 | **1.817e-05** |
| Sel:Trt | 66.62 | 66.62 | 1 | 8.0000 | 32.029 | **0.0004762** |

|  | npars | logLik | AIC | LRT | Df | Pr(>Chisq) |
| --- | --- | --- | --- | --- | --- | --- |
| <none> | 8 | -43.035 | 102.07 |  |  |  |
| (1 \| Block) | 7 | -43.037 | 100.07 | 0.0044 | 1 | 0.947229 |
| (1 \| Block:Selection) | 7 | -43.035 | 100.07 | 0.0000 | 1 | 0.999801 |
| (1 \| Block:Treatment) | 7 | -48.267 | 110.53 | 10.464 | 1 | **0.001217** |

*p*-values in bold are statistically significant. pairs(emmeans(fit\_ev\_hs\_24-30 h, ~Selection\*Treatment))
| contrast | estimate | SE | df | t.ratio | p.value |
| --- | --- | --- | --- | --- | --- |
| FCB heat shock - FSB heat shock | -7.2 | 0.912 | 8.00 | -7.893 | **0.0002** |
| FCB heat shock - FCB no shock | -65.5 | 2.669 | 4.49 | -24.541 | **<.0001** |
| FCB heat shock - FSB no shock | -65.4 | 2.669 | 4.49 | -24.504 | **<.0001** |
| FSB heat shock - FCB no shock | -58.3 | 2.669 | 4.49 | -21.844 | **<.0001** |
| FSB heat shock - FSB no shock | -58.2 | 2.669 | 4.49 | -21.806 | **<.0001** |
| FCB no shock - FSB no shock | 0.1 | 0.912 | 8.00 | 0.110 | 0.9995 |

*p*-values in bold are statistically significant. Degrees-of-freedom method: kenward-roger P value adjustment: tukey method for comparing a family of 4 estimates

**Table S2C.**
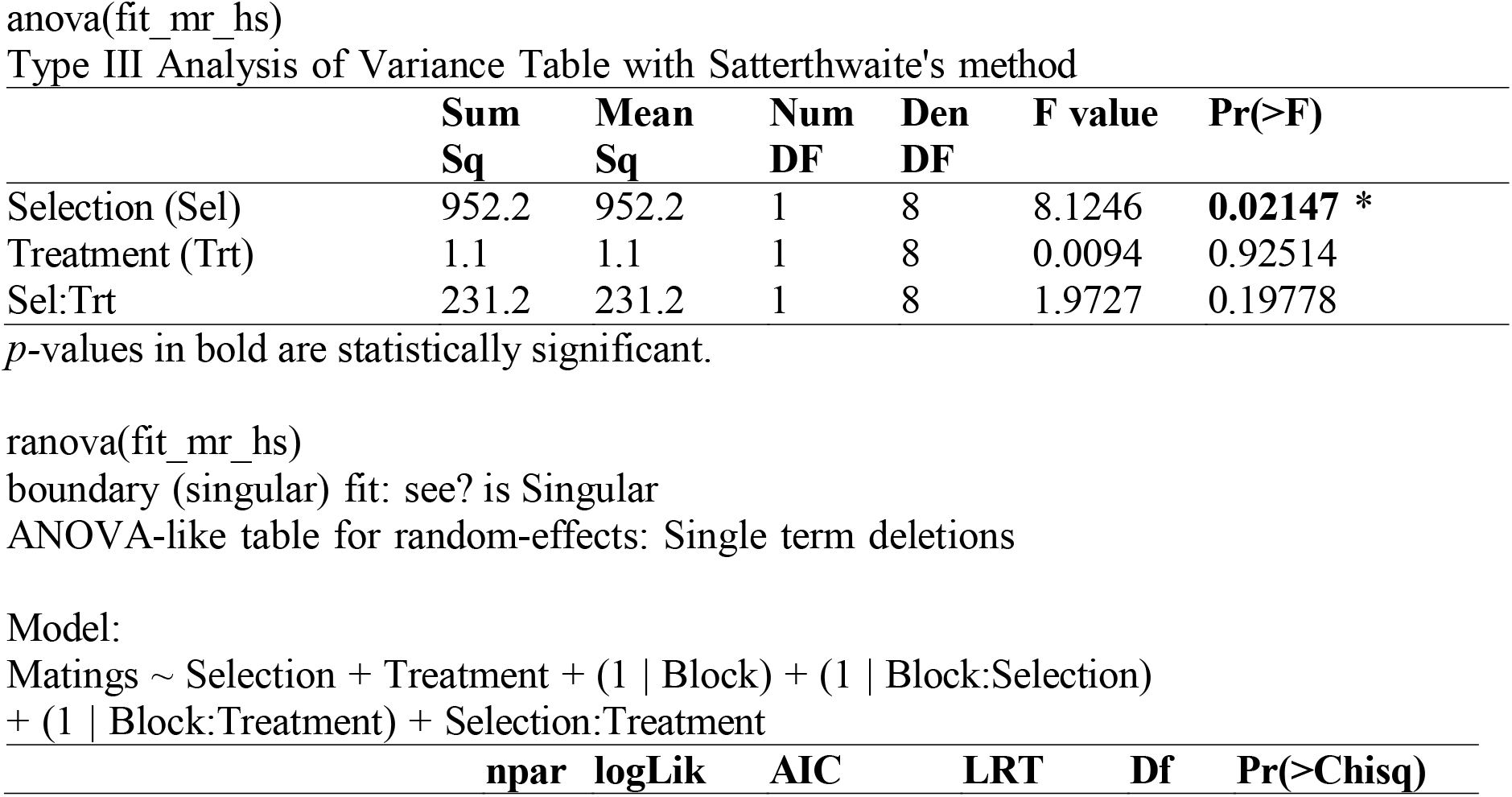
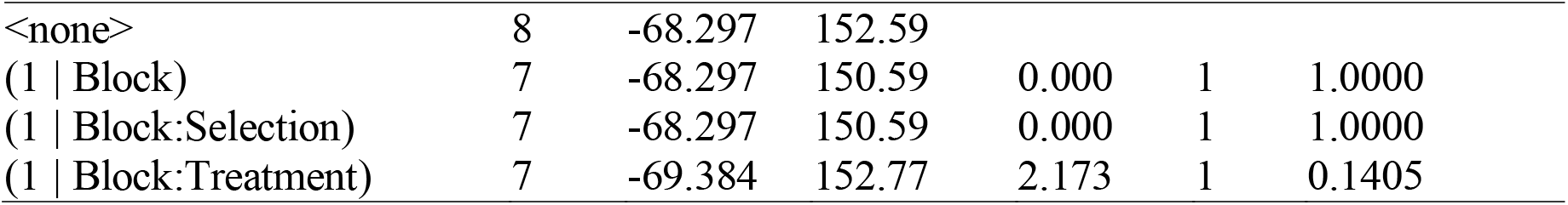
Increased mating frequency in the selected populations.

948 anova(fit\_mr\_hs) 949 Type III Analysis of Variance Table with Satterthwaite's method
|  | Sum Sq | Mean Sq | Num DF | Den DF | F value | Pr(>F) |
| --- | --- | --- | --- | --- | --- | --- |
| Selection (Sel) | 952.2 | 952.2 | 1 | 8 | 8.1246 | **0.02147** * |
| Treatment (Trt) | 1.1 | 1.1 | 1 | 8 | 0.0094 | 0.92514 |
| Sel:Trt | 231.2 | 231.2 | 1 | 8 | 1.9727 | 0.19778 |

|  | npars | logLik | AIC | LRT | Df | Pr(>Chisq) |
| --- | --- | --- | --- | --- | --- | --- |
| <none> | 8 | -68.297 | 152.59 |  |  |  |
| (1 \| Block) | 7 | -68.297 | 150.59 | 0.000 | 1 | 1.0000 |
| (1 \| Block:Selection) | 7 | -68.297 | 150.59 | 0.000 | 1 | 1.0000 |
| (1 \| Block:Treatment) | 7 | -69.384 | 152.77 | 2.173 | 1 | 0.1405 |

**Table S3.** Summary of the results from a two-factor mixed model ANOVA on adult survival in male (**S3A**) and female (**S3B**) post cold shock. And results from a two-factor mixed model ANOVA and on adult survival in male (**S3C**) and female (**S3D**) post heat shock employing selection regime (FSB and FCB) as the fixed factor crossed with blocks (1-5) as a random factor. *p-*values in bold are statistically significant.

| Trait | Effect | SS | MS<br>Num | DF<br>Num | DF<br>Den | *F* ratio | *P* |
| --- | --- | --- | --- | --- | --- | --- | --- |
| (**S3A**) | Selection (Sel) | 0.566 | 0.566 | 1 | 4 | 40.209 | 0.003 |
| Males | Block (Blk) | 0.296 | 0.074 | 4 | 4 | 5.245 | 0.069 |
| cold shock | Sel×Blk | 0.056 | 0.014 | 4 | 20 | 2.138 | 0.114 |
| (**S3B**) | Selection (Sel) | 0.637 | 0.637 | 1 | 4 | 52.076 | 0.002 |
| Females | Block (Blk) | 0.268 | 0.067 | 4 | 4 | 5.486 | 0.064 |
| cold shock | Sel×Blk | 0.049 | 0.012 | 4 | 20 | 2.240 | 0.101 |
| (**S3C**) | Selection (Sel) | 0.154 | 0.154 | 1 | 4 | 69.272 | 0.001 |
| Males | Block (Blk) | 0.227 | 0.057 | 4 | 4 | 25.460 | 0.004 |
| heat shock | Sel×Blk | 0.009 | 0.002 | 4 | 20 | 0.206 | 0.932 |
| (**S3D**) | Selection (Sel) | 0.105 | 0.105 | 1 | 4 | 60.146 | 0.001 |
| females | Block (Blk) | 0.026 | 0.006 | 4 | 4 | 3.737 | 0.115 |
| heat shock | Sel×Blk | 0.007 | 0.002 | 4 | 20 | 0.630 | 0.647 |

**Table S4.** Estimates and 95% confidence intervals (CIs) for relative hazard rates (i. e. the exponent of the coefficients) corresponding to various fixed parameters of the Cox’s proportional hazards model for **survivorship post infection**. Hazard rates are expressed relative to the default level of that fixed factor. The default level for selection regime is FCB, while the default level for sex is female. Hazard rates significantly greater than 1 correspond to poorer survivorship relative to the default level.

| Survivorship post infection |  |  |  |
| --- | --- | --- | --- |
| Fixed factors |  |  |  |
| Coefficient | Hazard ratio | Lower CL | Upper CL |
| SelectionFSB | 0.8149 | 0.6604 | 1.0056 |
| SexMale | 0.9528 | 0.6963 | 1.3037 |
| SelectionFSB:SexMale | 1.0309 | 0.7578 | 1.4025 |
| Random factors |  |  |  |
| Group | Variance |  |  |
| Block/Sex/Selection | 0.0013 |  |  |
| Block/Sex | 0.0343 |  |  |
| Block | <0.0001 |  |  |

**Table S5.** Estimates and 95% confidence intervals (CIs) for relative hazard rates (i. e. the exponent of the coefficients) corresponding to various fixed parameters of the Cox’s proportional hazards model for desiccation resistance of males (A) and females (B). Hazard rates are expressed relative to the default level of that fixed factor. The default level for selection regime is FCB. Hazard rates significantly greater than 1 correspond to poorer survivorship relative to the default level.

| (S5A) Male desiccation resistance |  |  |  |
| --- | --- | --- | --- |
| Fixed factors |  |  |  |
| Coefficient | Hazard ratio | Lower CL | Upper CL |
| SelectionFSB | 1.1600 | 0.3261 | 4.1262 |
| Random factors |  |  |  |
| Group | Variance |  |  |
| Block/Selection | 1.0324 |  |  |
| Block | 0.0714 |  |  |
| (S5B) Female desiccation resistance |  |  |  |
| Fixed factors |  |  |  |
| Coefficient | Hazard ratio | Lower CL | Upper CL |
| SelectionFSB | 0.5393 | 0.2760 | 1.0539 |
| Random factors |  |  |  |
| Group | Variance |  |  |
| Block/Selection | 0.2770 |  |  |
| Block | 0.2889 |  |  |

**Table S6.**
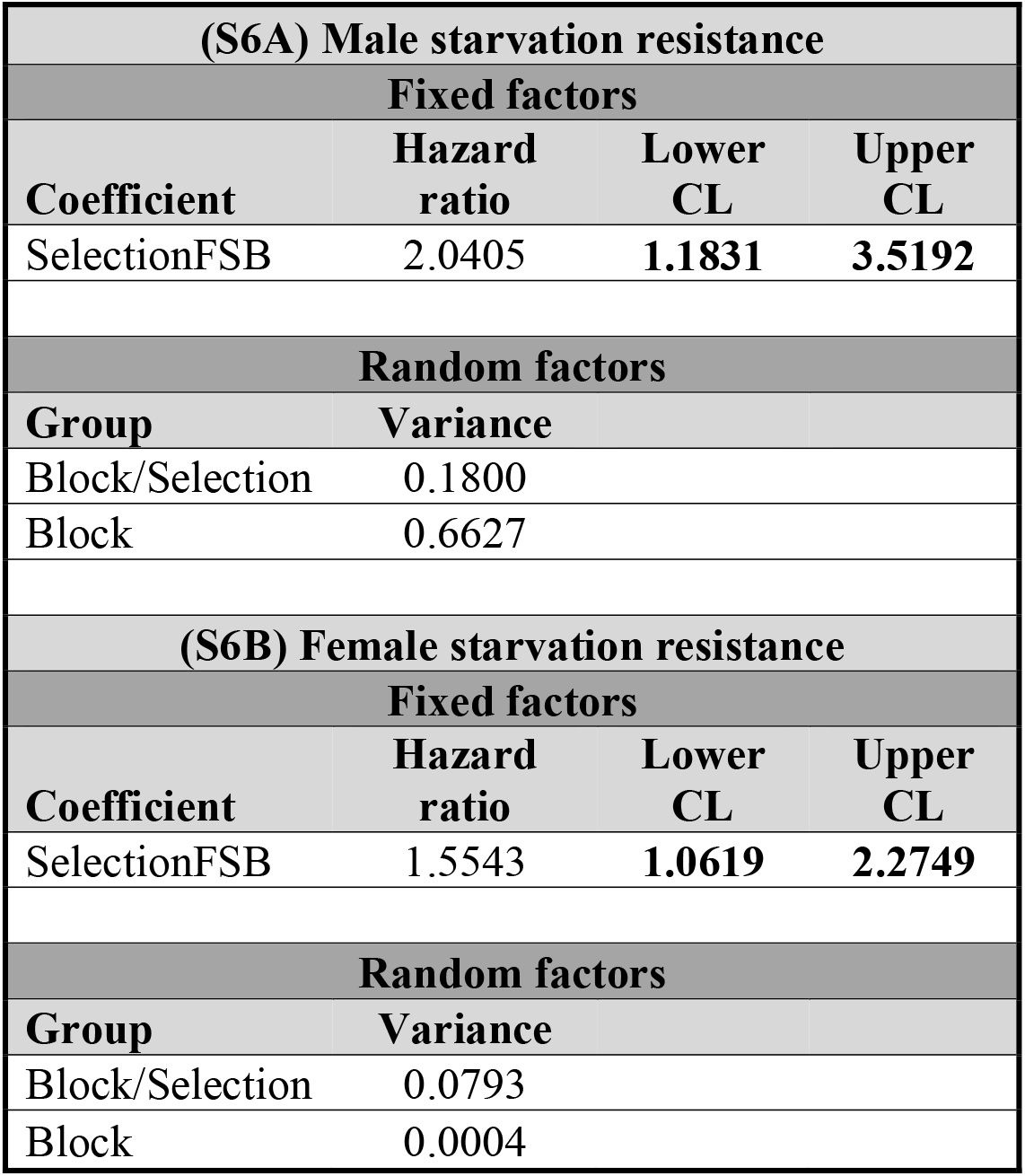
Estimates and 95% confidence intervals (CIs) for relative hazard rates (i. e. the exponent of the coefficients) corresponding to various fixed parameters of the Cox’s proportional hazards model for starvation resistance of males (A) and females (B). Hazard rates are expressed relative to the default level of that fixed factor. The default level for selection regime is FCB. Hazard rates significantly greater than 1 correspond to poorer survivorship relative to the default level.

| (S6A) Male starvation resistance |  |  |  |
| --- | --- | --- | --- |
| Fixed factors |  |  |  |
| Coefficient | Hazard ratio | Lower CL | Upper CL |
| SelectionFSB | 2.0405 | 1.1831 | 3.5192 |
| Random factors |  |  |  |
| Group | Variance |  |  |
| Block/Selection | 0.1800 |  |  |
| Block | 0.6627 |  |  |
| (S6B) Female starvation resistance |  |  |  |
| Fixed factors |  |  |  |
| Coefficient | Hazard ratio | Lower CL | Upper CL |
| SelectionFSB | 1.5543 | 1.0619 | 2.2749 |
| Random factors |  |  |  |
| Group | Variance |  |  |
| Block/Selection | 0.0793 |  |  |
| Block | 0.0004 |  |  |

